# Selective activation of IRE-1 safeguards restoration of translation and organismal rejuvenation from adult reproductive diapause

**DOI:** 10.64898/2026.03.18.712261

**Authors:** Qiang Fan, Ao Guo, Shujuan Wang, Wuling Yang, Yong-Hong Yan, Meng-Qiu Dong

## Abstract

In *C. elegans*, prolonged food deprivation during early larval development results in aging-like phenotypes that can be fully reversed upon refeeding, but it remains unknown whether and how this ability persists into later life stages. Here we subjected *C. elegans* of the last larval stage to starvation, driving them into adult reproductive diapause (ARD). During starvation, ARD worms exhibited a wide spectrum of aging-like phenotypes, including transcriptomic reprogramming accompanying cellular and functional declines. These phenotypes are largely restored within one day of refeeding, suggesting a rejuvenation effect. Time-series transcriptomics, proteomics, and follow-up analyses of the refeeding/rejuvenation process uncovered an intricate coordination between: (1) activation of the IRE-1 branch of UPR^ER^ (unfolded protein response of endoplasmic reticulum) to induce chaperone expression; (2) quiescence of the PEK-1 branch of UPR^ER^ to avoid translation suppression; (3) up-regulation of the translation machinery to boost protein synthesis. IRE-1, together with its downstream effector, XBP-1, play an essential role in boosting protein synthesis, which is required for complete rejuvenation from the ARD state. These findings indicate that coordination between a high protein synthesis activity and a high protein folding capacity is key to refeeding-associated rejuvenation.

**Highlights:**

- Refeeding rapidly reverses aging-like phenotypes in adult reproductive diapause
- Refeeding selectively activates the IRE-1 branch of UPR^ER^
- Refeeding restores protein synthesis in an IRE-1 dependent manner
- Rejuvenation requires enhanced translation activity and protein folding capacity

## Introduction

Aging is accompanied by phenotypic changes at all levels, from the invisible (e.g., various molecular damages) to the obvious (e.g., impaired sensory function and loss of muscle mass) ^1,2^. Targeting specific genes or pathways can restore these functions individually ^3–9^, for example, inhibiting prostaglandin-degrading enzyme 15-PGDH rejuvenates aged muscle ^9^. Yet, systemic reversal of aging phenotypes remains elusive and represents an ultimate goal of aging research.

In this regard, *Caenorhabditis elegans* provides a potential biological system to explore naturally occurring rejuvenation process. In first-stage (L1) *C. elegans* larvae, food deprivation leads to developmental arrest, with prolonged starvation leading to progressive accumulation of some aging-like phenotypes, including decline of mobility, fragmentation of mitochondrion, elevation of ROS and accumulation of protein aggregates ^10^. Yet, upon refeeding, worms can resume development and reverse the above aging-like phenotypes except the presence of protein aggregation ^10^. Further, worms growing out of the L1 arrest have a normal adult lifespan, suggesting that these worms may achieve complete functional rejuvenation ^10,11^. Alternatively, starvation of L2 or L4 larvae respectively leads to arrest in dauer (alternative L3) or adult reproductive diapause (ARD) states. Likewise, upon exit of dauer or ARD, worms display a normal adult lifespan, suggesting the occurrence of systemic rejuvenation ^12–16^.

It has been widely noted that juvenile exhibit stronger ability to repair tissue damage than adults ^17–19^. A unique aspect of ARD is that the worms enter the adult stage within one day after food deprivation. In other words, unlike L1 or dauer larvae, ARD worms rejuvenate, if occurs, as adults when their ability to repair damage is supposedly lower than larvae. Here, we investigated whether and how rejuvenation occurs in ARD worms. We found that ARD worms exhibit multi-level aging-like phenotypes during starvation, which are largely reversed within one day of refeeding. Functional assays coupled with transcriptomic and proteomic analyses in multiple worm models collectively revealed that IRE-1-dependent unfolded protein response of endoplasmic reticulum (UPR^ER^) coordinates restoration of protein translation activity and protein folding capacity to reverse starvation-induced aging-like phenotypes upon refeeding. Since ARD rejuvenation occurs during the adult stage, these findings provide new insights into how rebalancing proteostasis could potentially ameliorate structural and functional declines associated with natural aging.

## Results

### ARD worms exhibited multi-level aging-like phenotypes, which are reversed after refeeding

If deprived of food, *C. elegans* of the fourth−which is also the last−larval stage (L4), will complete development of the soma but suspend reproduction, entering a state called adult reproductive diapause (ARD) ^14^. If refed, ARD worms can regrow a full-sized gonad, produce progeny, and live−judging by their post-refeeding lifespan−as if they had never been starved ^14,15^. After verifying that ARD worms recovered from 15 or 5 days of complete starvation exhibited comparable adult lifespan to those of well-fed worms (Fig 1A), we next examined whether ARD worms develop aging-like phenotypes during starvation, and whether they reverse such phenotypes after refeeding. To this end, we treated mid-L4 stage *C. elegans* with complete starvation for up to 10 days (i.e., starved), for comparison with control worms without starvation treatment (i.e., well-fed) and worms starved for 10 days then refed for up to 10 days (i.e., refed; Fig 1B).

**Fig 1.**
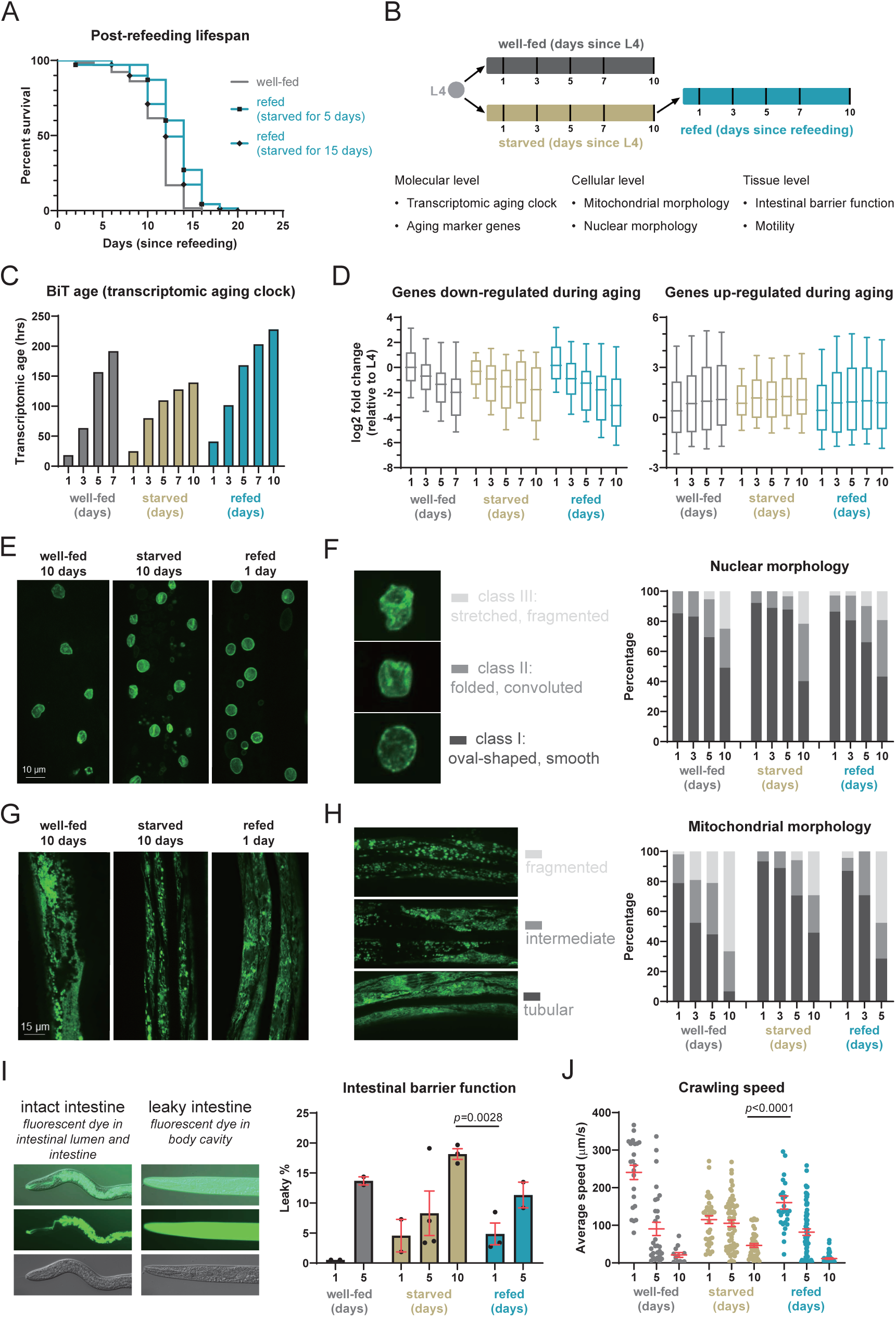
ARD worms exhibited multi-level aging-like phenotypes, which were reversed within 1 day of refeeding. (A)Prolonged starvation did not shorten post-feeding lifespan in ARD worms. (B)Experimental design of characterizing aging-like phenotypes in ARD worms. (C)Transcriptomic age estimated by BiT-age. (D)mRNA abundance of aging marker genes defined in ^14^, including 67 down-regulated genes and 260 up-regulated genes. (E and F) Hypodermal nuclear morphology. EMR-1::GFP labeled inner nuclear membrane. n>30 animals per timepoint. (G and H) Body wall muscle mitochondrial morphology. GFP localized in mitochondrial matrix. n>30 animals per timepoint. Intestinal barrier function. Leaky percentage was calculated as the proportion of worms exhibiting leakage of fluorescein sodium through the intestinal barrier into the body cavity. Each dot represents one independent biological replicate. n>150 animals per timepoint. *p*, two-tailed unpaired t-test. (J)Crawling speed. Each dot represents one worm. *p*, unpaired Mann-Whitney U test. Error bars are means + SEM.

Firstly, we sought to determine how transcriptomic signatures of aging change during starvation and refeeding. BiT age is a transcriptome-based aging clock that can capture the transcriptomic signatures of aging ^20^. We found that transcriptomic age increased in both starved and normally aging well-fed worms; after refeeding, transcriptomic age was reset within one day and then progressed as in the well-fed group (Fig 1C). Analysis of aging-related marker expression ^21^ revealed that these genes showed the same transcriptional trajectory between starved and well-fed groups, whereas they reversed to a pre-starvation signature in the refed group, then subsequently commenced with an aging-aligned trajectory (Fig 1D). For example, down-regulated genes decreased in transcription during both normal aging and starvation, but showed fully restored transcript levels within one day of refeeding before progressively declining (Fig 1D). These results suggested that starvation induces transcriptomic signatures of aging, while refeeding completely abrogates these signatures.

We then examined subcellular changes in nuclear morphology and mitochondrial morphology. Abnormalities in nuclear morphology arise during aging ^22^. We monitored changes in nuclear morphology in hypodermis tissue by labeling inner nuclear membrane with EMR-1::GFP. Most nuclei were oval-shaped at day 1 of starvation (class I); however, the percentage of nuclei with a shriveled or wrinkled morphology (class II, class III) gradually increased by day 10 of starvation (Fig 1E-F). In contrast, oval nuclei were restored within one day of refeeding, then again gradually shifted toward the aged nuclear phenotype (Fig 1E-F). Mitochondria in body wall muscle cells form tubular neywork in young worms, but fragment during aging (Fig 1G-H) ^23^. To visualize mitochondrial morphology, we performed confocal imaging on worms expressing matrix-targeted GFP. We observed that starved worms indeed showed mitochondrial fragmentation, whereas tubular cristae were completely restored after refeeding, then gradually fragmented as in well-fed controls (Fig 1G-H). These results thus indicated that refeeding induces restoration of nuclear morphology in hypodermis as well as mitochondrial networks in body wall muscle cells.

To examine tissue-level physiological changes, we focused on intestinal barrier function and motility. In well-fed controls administered with the fluorescein sodium, we observed that the proportion of worms exhibiting leakage through the intestinal barrier into the body cavity significantly increased with age, indicating progressive weakening of intestinal barrier function (Fig 1I) ^24^. The proportion of worms with compromised integrity of the intestinal barrier was also increased by 10 days of starvation. In contrast, this leakage was largely attenuated within one day of refeeding (Fig 1I). Additionally, motility analysis also showed a decline with normal aging (Fig 1J), which has been previously established as a conserved phenotype of biological aging ^25^. While starved worms also exhibited decreased crawling speed, we found that refed worms crawled significantly faster (Fig 1J), indicating that refeeding leads to restoration of intestinal and muscular function.

Together, these observations indicated that starvation-induced ARD worms exhibit multi-level deterioration, phenotypically resembling aged worms at a comparable progression rate; yet, refeeding can rejuvenate ARD worms within one day before re-initiating the aging process.

### Refeeding-associated rejuvenation initiates within several hours

In light of our above results showing that rejuvenation occurs within one day of refeeding, we next sought to characterize the temporal dynamics of the rejuvenation process at finer resolution. Further BiT age analysis showed that transcriptomic age slightly decreased at 3 hours post-refeeding, then significantly decreased at 6 hours, reaching its lowest value at 18 hours (Fig 2A). Similarly, transcript abundance of aging marker genes started to revert by 3 hours of refeeding, and reached a plateau by 18 hours. (Fig 2B). The proportions of oval nuclei and tubular mitochondria increased by 6 hours of refeeding, and this increasement proceeded until 24 hours of refeeding (Fig 2C-D). Additionally, the proportion of worms with intestinal leakage slightly decreased after 6 hours of refeeding, but remained relatively high through 12 hours, suggesting that intestinal barrier function was predominantly restored between 12 and 24 hours of refeeding (Fig 2E). Similarly, crawling speed did not change within 6 hours of refeeding, and only partially increased after 12 hours of refeeding (Fig 2F). Overall, these results indicated that the rejuvenation process initiates within several hours of refeeding, and completes within one day.

**Fig 2.**
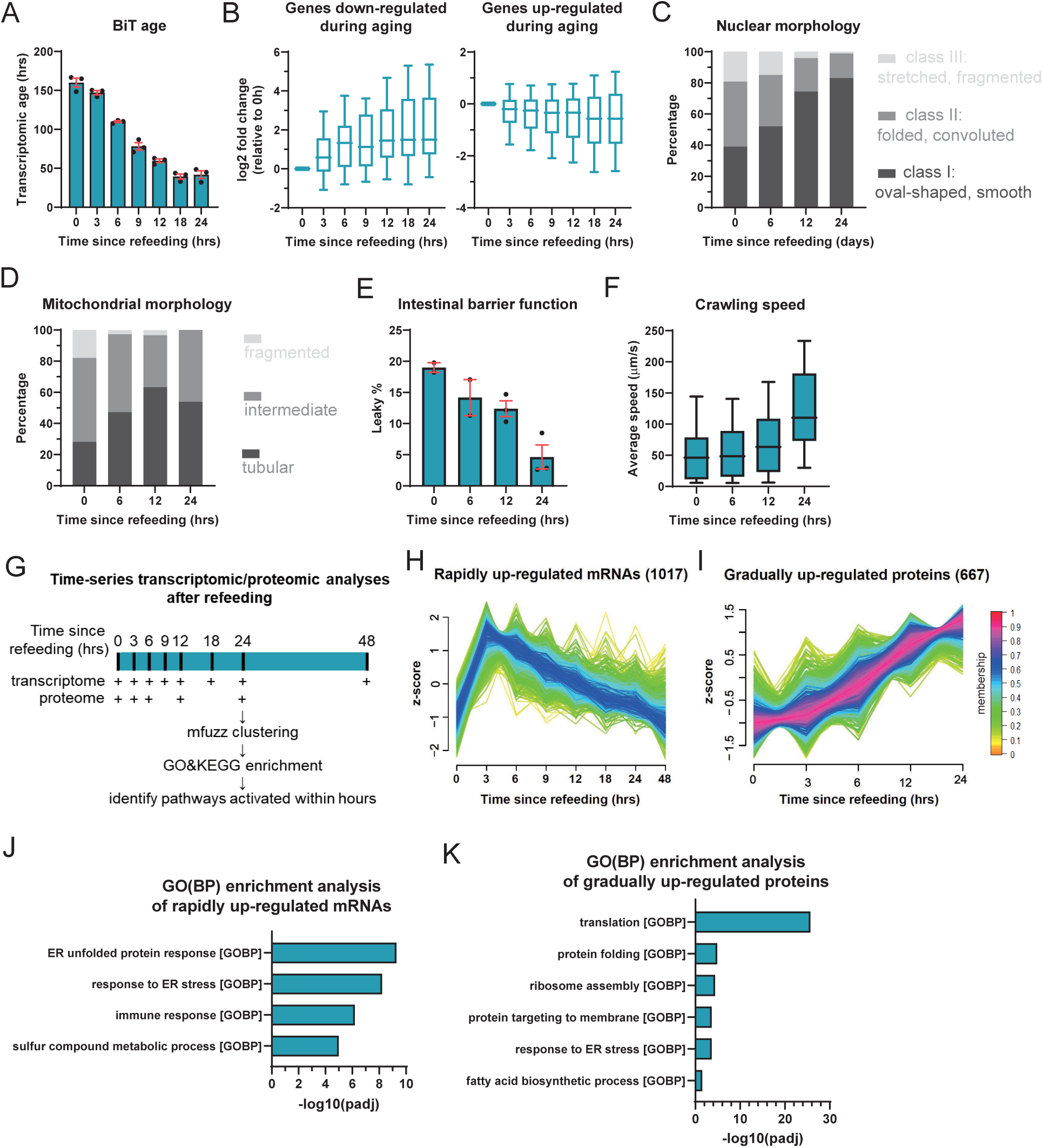
Refeeding-associated rejuvenation initiates within several hours. (A)Transcriptomic age estimated by BiT-age. Each dot represents one independent biological replicate. (B)mRNA abundance of aging marker genes defined in ^14^. (C)Hypodermal nuclear morphology. EMR-1::GFP labeled inner nuclear membrane. n>30 animals per timepoint. (D)Body wall muscle mitochondrial morphology. GFP localized in mitochondrial matrix. n>30 animals per timepoint. (E)Intestinal barrier function. Each dot represents one independent biological replicate. n>150 animals per timepoint. (F)Crawling speed. n>100 animals per timepoint. (G)Experimental design of time-series transcriptomic/proteomic analyses of the refeeding process. (H and I) Temporal expression pattern of rapidly up-regulated mRNAs (H) and gradually up-regulated proteins (I). (J and K) GO(BP) enrichment analyses of rapidly up-regulated mRNAs (J) and gradually up-regulated proteins (K). Error bars are means + SEM.

### Refeeding rapidly activates UPR^ER^ and protein synthesis

The refeeding-associated rejuvenation initiated within several hours, implying that the underlying regulatory mechanism was also engaged on the same rapid timescale. Thus, we conducted a time course transcriptomic and proteomic analyses to identify pathways that are rapidly activated by refeeding (Fig 2G).

To assess the global transcriptomic shifts during refeeding, we performed principal component analysis (PCA). Notably, refed 3h group speedily departed from starvation group, and the following timepoints migrated much slower (Fig S1B). To estimate the rate of transcriptomic remodeling, we divided the number of differentially expressed genes (DEGs) by the interval of adjacent time points (i.e., DEG generation rate). As expected, the DEG generation rate was highest in the first 3 hours of refeeding, then markedly decreased and remained low for the duration of the observation period (Fig S1C), suggesting that the transcriptomic response to refeeding was largely established within 3∼6 hours. Subsequent mfuzz clustering analysis according to temporal expression patterns ^26^ identified a cluster comprised genes that were rapidly and transiently activated after refeeding: transcript abundance shot up within 3 hours, following by a gradual decline (Fig 2H; cluster 4 in Fig S1D). Gene ontology (GO) enrichment analyses showed that these genes were significantly enriched in multiple homeostasis-regulating biological processes (Fig 2J), including most prominently unfolded protein response of endoplasmic reticulum (UPR^ER^).

In contrast to the rapid shifts observed in the transcriptome, PCA of the proteomic data revealed a more progressive reshaping of the proteome, with samples from each timepoint evenly positioned along the recovery trajectory (Fig S2B). Consistently, dividing the number differentially expressed proteins (DEPs) by interval, to estimate a DEP generation rate between adjacent time points, showed that DEP generation rate remained steady until 12 hours of refeeding and slightly decreased thereafter (Fig S2C). Subsequent clustering and enrichment analyses indicated that proteins involved in UPR^ER^ or translation are overrepresented in a specific group of proteins (Fig 2K), which gradually accumulated after refeeding (Fig 2I; cluster 2 in Fig S2D). Additionally, we noted that UPR^ER^- and translation-related proteins had markedly lower abundance in starved worms compared to well-fed L4 worms (Fig S2E-F), suggesting these proteins responsible for ER protein folding capacity and translation activity were depleted during starvation and restored upon refeeding.

Thus, refeeding rapidly activates diverse cellular programs involved in homeostasis, including UPR^ER^ and protein synthesis.

### Refeeding activates IRE-1-XBP-1 axis to up-regulate UPR^ER^ genes

To further investigate a possible role of UPR^ER^ in rescuing aging-related phenotypes after starvation, we sought to reveal how refeeding activates UPR^ER^. Comparing our transcriptomic and proteomic datasets, we found that refeeding led to activation of UPR^ER^ genes at both the mRNA and protein levels (Fig 3A). Yet, while protein levels accumulated progressively, mRNA expression for ∼66% of these genes peaked early and then moderated, whereas the remaining ∼30% maintained steady induction (Fig 3A). *hsp-4* is a canonical marker gene for the UPR^ER^, and encodes the *C. elegans* homolog of BiP/GRP78 ^27^. Using a transcriptional reporter strain expressing GFP driven by the *hsp-4* promoter, we found that after refeeding, GFP fluorescence gradually increased until 12 hours and declined thereafter (Fig 3B). Using a HSP-4::mNeonGreen knock-in strain, we found that the fluorescence intensity gradually increased until 24 hours after refeeding (Fig 3C). These observations validate our transcriptomic and proteomic findings.

**Fig 3.**
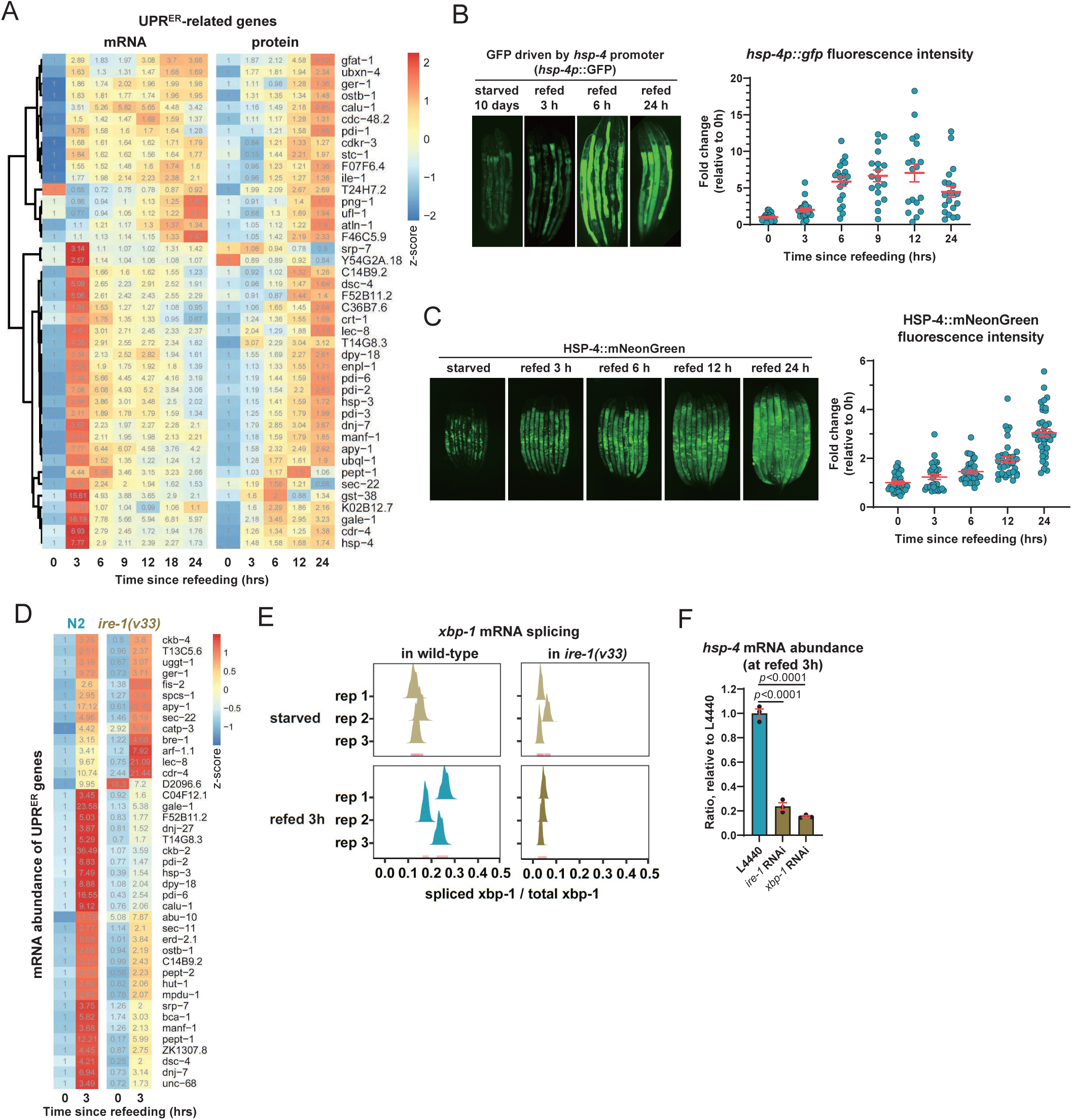
Refeeding rapidly and transiently activates IRE-1-mediated UPR^ER^. (A)mRNA and protein abundance of UPR^ER^ genes after refeeding. The color scale indicates z-score, and numbers labeled within each cell represent fold change relative to 0 h. (B)Rapid and transient activation of hsp-4 promoter after refeeding. Each dot represents one animal. (C)Fluorescence intensity of HSP-4::mNeonGreen gradually increased after refeeding. Each dot represents one animal. (D)Transcriptional upregulation of UPR^ER^ genes was determined by *ire-1*. The color scale indicates z-score, and numbers labeled within each cell represent fold change relative to N2 refed 0 h. (E)Splicing of xbp-1 mRNA increased in refed wild-type N2 worm but not in refed *ire-1(v33)* worms. (F)qPCR analysis showed that up-regulation of *hsp-4* was inhibited by knocking down *xbp-1*. Each dot represents one independent biological replicate. *p*, two-tailed unpaired t-test. Error bars are means + SEM.

The ribonuclease inositol-requiring protein-1 (IRE-1) is a major regulator of canonical UPR^ER 28^. Upon ER stress, activated IRE-1 catalyzes the unconventional splicing of *xbp-1* (X-box binding protein-1) mRNA. The resulting spliced isoform (*xbp-1s*) encodes a transcription factor that upregulates a suite of UPR genes—including the heat-shock protein 4 (*hsp-4)*—thereby enhancing protein folding and degradation capacity ^28^.

To test whether IRE-1 was required for UPR^ER^ pathway activation in response to refeeding, we conducted RNA-seq in refed *ire-1(v33)* mutant and refed wild-type N2 worms. This analysis revealed that upregulation of most UPR^ER^ genes upon refeeding was markedly attenuated in *ire-1(v33)* worms compared to wild-type controls (Fig 3D). Moreover, alternative splicing detection with rMATS (replicate multivariate analysis of transcript splicing ^29^) indicated that splicing of *xbp-1* mRNA increased upon refeeding in wild-type worms, but not in the *ire-1(v33)* mutant (Fig 3E). Further, we observed that *hsp-4* upregulation at 3h of refeeding was profoundly inhibited by RNAi-mediated *ire-1* or *xbp-1* knockdown (Fig 3F). These results thus indicated that the IRE-1-XBP-1 axis was required for upregulation of UPR^ER^ genes following refeeding in starved worms.

### Refeeding-associated rejuvenation relies on IRE-1-mediated UPR^ER^

To next investigate how IRE-1-dependent UPR^ER^ activation contributes to rejuvenating starved worms after refeeding, we conducted BiT analysis in the *ire-1(v33)* strain. Notably, transcriptomic age decreased slowly in *ire-1(v33)* worms compared to that of wild-type N2 following refeeding (Fig 4A). In line with these results, we found that the *ire-1 (v33)* mutant retained abnormalities in hypodermal nuclear morphology after refeeding (Fig 4B).

**Fig 4.**
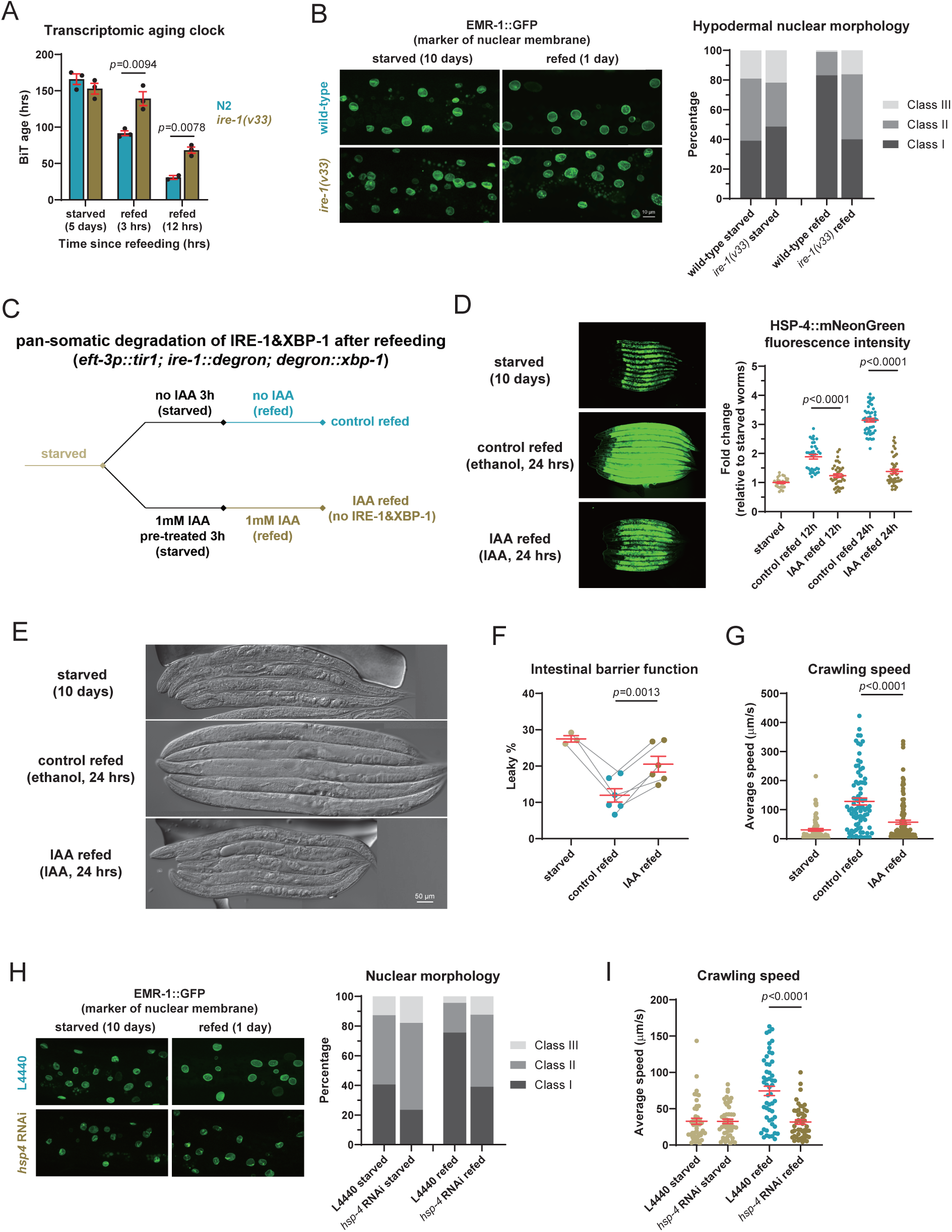
IRE-1-mediated UPR^ER^ promoted refeeding-associated rejuvenation. (A)In refed *ire-1(v33)*, transcriptomic age was reversed slowly. Each dot represents one independent biological replicate. *p*, two-tailed unpaired t-test. |(B)In refed *ire-1(v33)*, abnormal nuclear morphology was not reversed. n>30 animals per timepoint. (C)Experimental design of auxin-inducible degradation of IRE-1 and XBP-1 in refed worms. (D)Degradation of IRE-1 and XBP-1 profoundly inhibited accumulation of HSP-4 in refed worms. Each dot represents one animal. *p*, unpaired Mann-Whitney U test. (E, F and G) Degradation of IRE-1 and XBP-1 inhibited somatic regrowth (E), reversal of intestinal leaky (F) and restoration of crawling speed (G) in refed worms. In (F), each dot represents one independent biological replicate, n>150 animals per timepoint, *p* represents two-tailed unpaired t-test. In (G), each dot represents one animal, *p* represents unpaired Mann-Whitney U test. (H and I) Knocking down *hsp-4* inhibited restoration of nuclear morphology (H) and crawling ability (I). Bacteria contained empty L4440 plasmid served as control. In (H), n>30 animals per timepoint. In (I), each dot represents one animal, *p* represents unpaired Mann Whitney U test

To determine whether IRE-1-mediated UPR^ER^ is specifically required during the refeeding phase, rather than the starvation phase, to enable rejuvenation, we used an auxin-inducible degradation (AID) system to specifically deplete IRE-1 and/or XBP-1 after refeeding (Fig 4C) ^30^. While IRE-1 degradation largely suppressed transcriptional up-regulation of *hsp-4* (Fig S3A), HSP-4 protein accumulation was only partially suppressed (Fig S3B). XBP-1 degradation led to a relatively slight decrease in *hsp-4* upregulation in refed worms (Fig S3C). These results indicated that IRE-1 or XBP-1 degradation alone could not efficiently inhibit UPR^ER^. In contrast, simultaneously depleting IRE-1 and XBP-1 resulted in profound suppression of *hsp-4* upregulation (Fig S3D) and HSP-4 protein accumulation (Fig 4D). Using this double AID system, we found that whereas starved no-IAA control worms resumed growth, restored intestinal function, and regenerated germline cells upon refeeding, IAA-treated worms ingested food but failed to recover (Fig 4E and S4). Simultaneous depletion of IRE-1 and XBP-1 also blocked the rescue of intestinal leakage (Fig 4F) as well as restoration of locomotion (Fig 4G). Additionally, in worms expressing TIR1 but lacking degron tag, IAA treatment failed to suppress *hsp-4* up-regulation (Fig S5A), resumption of growth (Fig S5B), reversal of intestinal leakage (Fig S5C), or restoration of locomotion (Fig S5D) following refeeding. Together, these results demonstrated that the impaired rejuvenation phenotype of double AID worms was the result of IRE-1/XBP-1 depletion, rather than toxicity or off-target effects of IAA.

Finally, to determine whether the IRE-1-XBP-1 axis affects rejuvenation by modulating protein folding capacity, we induced *hsp-4* knockdown by RNAi. We found that *hsp-4* knockdown phenocopied the effects of IRE-1/XBP-1 depletion, as these worms showed no restoration of nuclear morphology (Fig 4H) or crawling ability compared to L4440 empty vector-treated control worms following refeeding (Fig 4I). Taken together, these results indicated that IRE-1-mediated UPR^ER^ activation is necessary for refeeding-associated rejuvenation.

### After refeeding, quiescence of PEK-1-mediated UPR^ER^ cooperates with translation reactivation

As canonical UPR^ER^ can also inhibit translation to reduce protein folding burden via PEK-1-mediated eIF2a phosphorylation, we also investigated whether the PEK-1-eIF2a branch participated in rejuvenation. After confirming by western blot that exposure to the thiol-reducing reagent dithiothreitol (DTT), which could induce ER stress, led to increased levels of Ser49-phosphorylated eIF2a in wild-type, but not *pek-1(ok275)* mutant worms (Fig S6), we examined whether eIF2a phosphorylation increased following refeeding. However, phosphorylated eIF2a gradually decreased in both wild-type controls (Fig 5A) and *pek-1(ok275)* mutant worms (Fig 5B) after refeeding. Thus, we concluded that the PEK-1 branch remained inactive after refeeding, possibly allowing enhanced translation.

**Fig 5.**
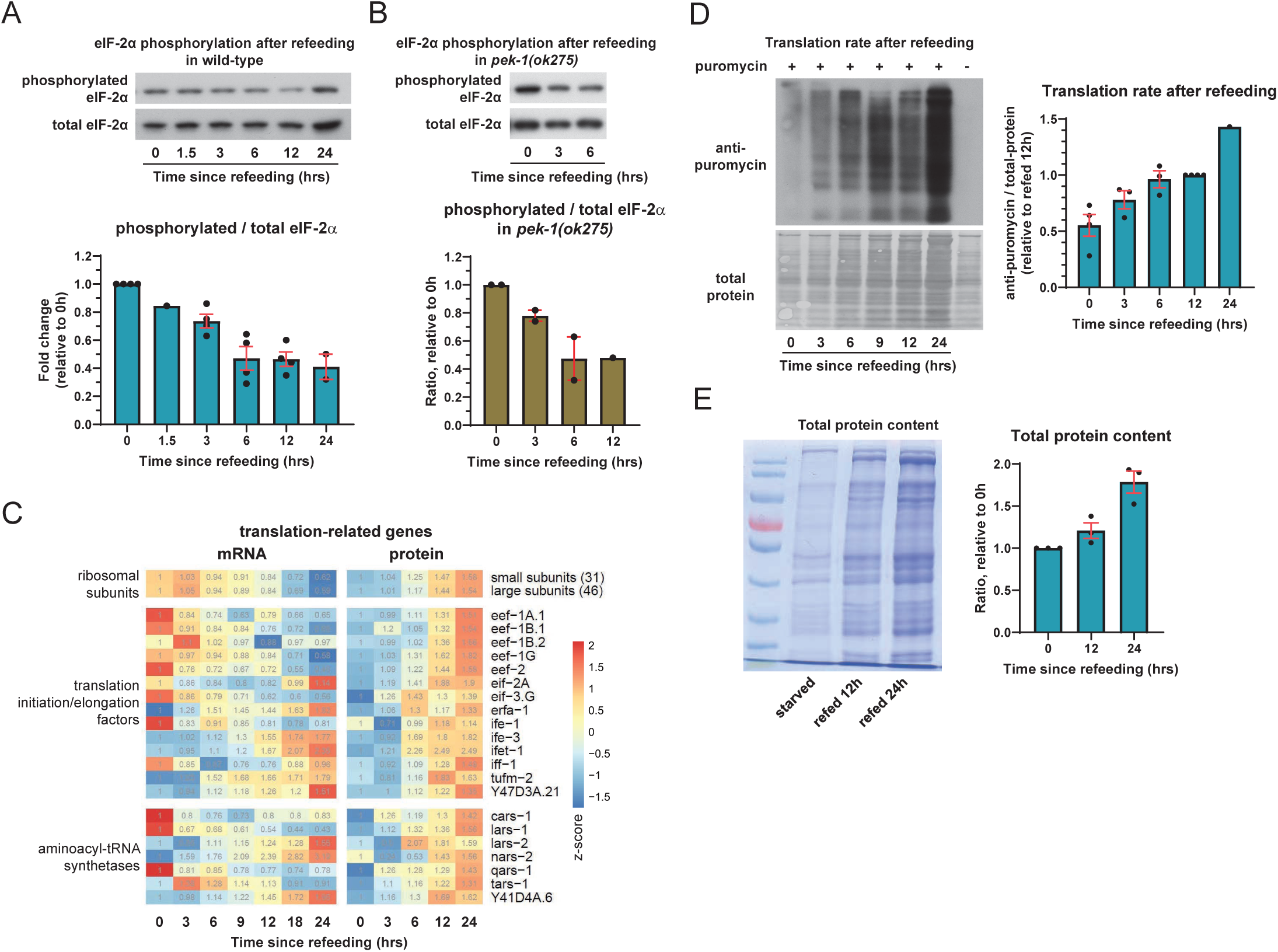
Quiescence of PEK-1-mediated UPR^ER^ cooperates with increased translation activity. (A and B) Phosphorylation of eIF2a decreased after refeeding in wild-type (A) and *pek-1(ok275)* (B). (C)mRNA and protein abundance of translation-related genes after refeeding. The color scale indicates z-score, and numbers labeled within each cell represent fold change relative to 0 h. (D)Translation activity (measured by SUnSET assay) increased after refeeding. (E)Total protein content increased after refeeding. n=200 animals per timepoint. Each dot represents one independent biological replicate. Error bars are means + SEM.

Revisiting our above proteomic analysis revealed accumulation of translation-related proteins in refed worms (Fig 2I and 2K). However, comparison with transcriptomic data showed no concomitant increase in transcription of these genes, suggesting post-transcriptional regulation might support this increased translational response to refeeding (Fig 5C). Since changes in the abundance of translation machinery can substantially affect translation activity, we measured translation rates during starvation and refeeding. Surface sensing of translation (SUnSET) assays ^31^ indicated that translation rates decreased during starvation (Fig S7A) and gradually increased after refeeding (Fig 5D), which led to accumulation of total protein (Fig 5E). These results suggested that refeeding coordinates proteostasis through three complementary mechanisms: augmentation of translation machinery proteins leads to increased translation activity, activation of IRE-1-XBP-1 axis enhance protein folding ability, quiescence of PEK-1-eIF2a axis avoids repression of translation.

### IRE-1-mediated UPR^ER^ supports translation reactivation and thereby reversal of aging-like phenotypes

To verify whether translation reactivation upon refeeding requires augmented protein folding capacity via activation of the IRE-1-mediated UPR^ER^, we assessed translation rates after refeeding in dual AID worms with IRE-1 and XBP-1 depletion. SUnSET assays indicated that refeeding-induced translation reactivation was indeed blocked in the absence of IRE-1 and XBP-1 (Fig 6A). This effect was confirmed by SDS-PAGE assays showing no increase in total protein contents after refeeding in dual AID worms compared to control worms without IRE-1/XBP-1 depletion (Fig 6B). In refed *ire-1(v33)* mutant worms, SUnSET assays and total protein extractions also showed no increase in translation activity (Fig S7B) or protein contents compared with wild-type N2 controls (Fig S7C), supporting the likelihood that IRE-1-mediated UPR^ER^ disruption blocked refeeding-induced translation reactivation, rather than toxicity artifacts of IAA exposure.

**Fig 6.**
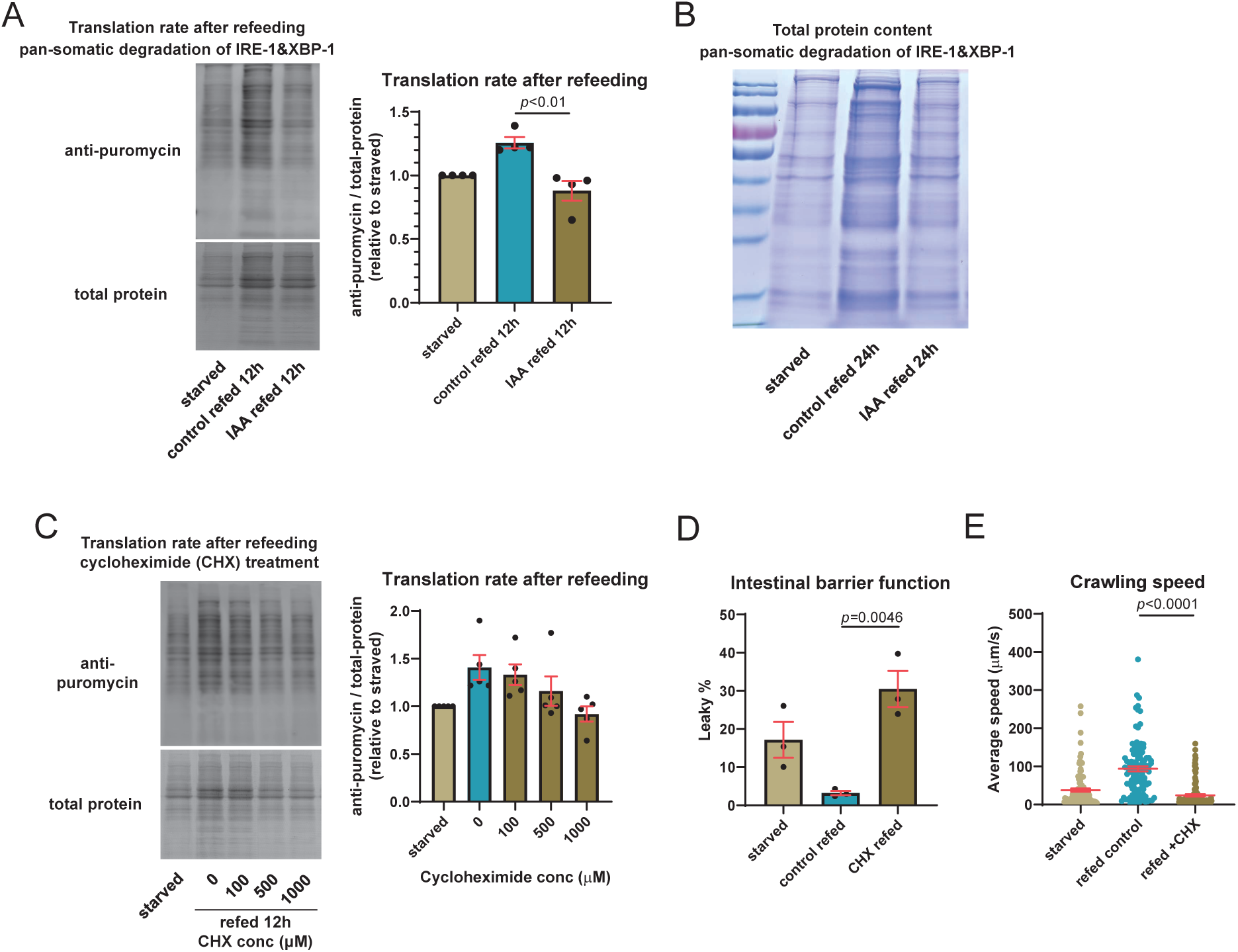
Selective activation of IRE-1-mediated UPR^ER^ supports efficient protein synthesis, which is required for reversal of aging phenotypes. (A)Degradation of IRE-1 and XBP-1 inhibited translation reactivation in refed worms. Each dot represents one independent biological replicate. *p*, two-tailed unpaired t-test. (B)Degradation of IRE-1 and XBP-1 inhibited accumulation of total protein in refed worms. (C)Cycloheximide treatment inhibited translation reactivation in refed worms. Each dot represents one independent biological replicate. (D and E) Cycloheximide treatment inhibited reversal of intestinal leaky (D) and restoration of crawling speed (E) in refed worms. In (D), each dot represents one independent biological replicate, n>150 animals per timepoint, *p* represents two-tailed unpaired t-test. In (E), each dot represents one animal, *p* represents unpaired Mann-Whitney U test. Error bars are means + SEM.

Finally, to verify that refeeding-associated rejuvenation requires IRE-1-dependent translational reactivation, we examined whether rejuvenation occurred under translational inhibition with cycloheximide (CHX), a widely used translation inhibitor. SUnSET assays revealed that 1000 µM CHX could profoundly attenuate the refeeding-induced increase in translation rates, but conferred negligible effect on translational reactivation at lower doses (e.g., 100 µM CHX; Fig 6C). Additionally, 1000 µM CHX treatment resulted in complete suppression of refeeding-induced reversal of intestinal leakage (Fig 6D) as well as restoration of locomotion (Fig 6E). These findings indicated that IRE-1-dependent translational reactivation is indeed required for rejuvenation of aging-like phenotypes.

## Discussion

Our study demonstrates that the capacity for systemic rejuvenation in *C. elegans* persists into the adult reproductive diapause (ARD) stage. We show that refeeding-associated rejuvenation is driven by a coordinated proteostatic program that selectively activates the IRE-1-XBP-1 axis to enhance folding capacity while bypassing PEK-1-mediated translation suppression. This uncoupling of UPR^ER^ branches sustains the robust protein synthesis necessary for rejuvenation. Collectively, these findings establish a mechanistic framework for how organisms recalibrate their proteostasis networks after severe metabolic stress.

Although reversal of some aging-like phenotypic defects has been observed in dauer and L1 larvae ^10,13,16^, it has remained uncertain whether the capacity for rejuvenation is maintained throughout the entire lifecycle. In the present study, we observed that aging-like alterations in transcriptomic program, organellar morphology, and tissue function were largely abrogated upon refeeding in starved ARD worms, suggesting that rejuvenation is not restricted to early larval stages, but rather persists at least through the transition to adult stage. Whether *C. elegans* retains the capacity for rejuvenation after starvation conditions at advanced ages remains an open question.

Results in this study demonstrate that protein synthesis and folding capacity both increase after refeeding. Specifically, we found that translation-related proteins increase in abundance upon refeeding, and this restoration is likely mediated via post-transcriptional regulation (Fig 5C). These findings align well with previous report that these proteins were selectively translated during dauer recovery ^32^. Our transcriptomic analyses showed that refeeding activates the IRE-1-XBP-1 branch of the UPR^ER^, which in turn transcriptionally up-regulates chaperone-encoding genes, while the PEK-1-eIF2a axis remained inactive. In classic ER stress response, both IRE-1 and PEK-1 are activated, respectively increasing the abundance of chaperones while reducing the unfolded protein load. However, under conditions such as refeeding, which requires efficient protein synthesis, activation of the PEK-1-eIF2a axis can inhibit translation, consequently impeding recovery from starvation (Fig 6D and 6E). We therefore propose that refeeding selectively activates IRE-1 to safeguard proteostasis without hindering protein synthesis. Refeeding has been reported to induce UPR^ER 33–40^ and promote translation ^41,42^ in other species. Further investigation is warranted in future studies to determine whether specifically activating the IRE-1-mediated UPR^ER^ is an evolutionarily conserved starvation recovery strategy.

The refeeding-associated rejuvenation initiated within several hours (Fig 2A-F). Our time-series transcriptomic and proteomic analysis thus lie a foundation for studies focusing on rejuvenation mechanism. Leveraging these analyses, we found that phenotypic restoration relies on the IRE-1-mediated UPR^ER^ to support efficient protein synthesis. Notably, refeeding-induced UPR^ER^ activation is not restricted to ARD worms, and can also be observed in L1 arrest (Fig S8A) ^10,43^ and dauer (Fig S8B) ^32^. Additionally, IRE-1 has been previously demonstrated as required for L1 recovery ^10^. These findings thus lead us to propose that IRE-1-dependent translational reactivation may serve as a rejuvenation mechanism across life stages in *C. elegans*.

Although we determined that refeeding activates the IRE-1-mediated UPR^ER^ (Fig 3), whether transcriptional up-regulation of UPR^ER^ genes fully relies on IRE-1-XBP-1 axis remains unclear. In AID worms, XBP-1 depletion partially inhibited *hsp-4* up-regulation upon refeeding (Fig S3C). Consistent with that finding, transcriptomic analyses indicated that *xbp-1* splicing is completely abolished in the *ire-1(v33)* mutant (Fig 3E), but UPR^ER^ gene up-regulation is only partially inhibited (Fig 3D). Additionally, we observed nuclear translocation of XBP-1::GFP following DTT treatment, but not in starved or refed worms (Fig S9), suggesting XBP-1 is only partially required, and therefore plays a relatively minor role, in up-regulating UPR^ER^ genes in response to refeeding. Although the *ire-1(v33)* mutant lacks the genomic region encoding the luminal domain, which senses misfolded proteins in ER lumen, this *v33* deletion may not impact the shorter mRNA isoform of *ire-1* ^44^. Transcriptional profiling showed that UPR^ER^ genes were still partially up-regulated upon refeeding in the *ire-1(v33)* mutant (Fig 3D), indicating that the deletion impaired, but did not block, IRE-1-mediated UPR^ER^ activation. Thus, we propose transcriptional activation of UPR^ER^ partially relies on luminal domain of IRE-1, or other IRE-1-independent regulators may exist. The three branches of the UPR^ER^ are known to act in overlapping pathways ^45^. However, *pek-1, atf-4, or atf-6* knockdown did not inhibit *hsp-4* up-regulation (Fig S10), suggesting that neither the PEK-1-ATF-4 axis nor ATF-6 engage in refeeding-associated UPR^ER^. Our current work thus reveals a role of IRE-1-XBP-1 axis in regulating refeeding-associated UPR^ER^, while also implicating the involvement of additional regulatory factors.

One compelling possibility is that increased translation leads to accumulation of misfolded proteins, which in turn activates IRE-1. However, this model struggles to explain why PEK-1, which also possesses misfolded protein sensing capability, is not activated. Also, as we discussed above, luminal domain of IRE-1 is not entirely required for UPR^ER^ activation. Further study is necessary to determine what role, if any, protein folding defects may play in IRE-1 activation. Alternatively, nutrient sensing pathways may modulate IRE-1 activity, as previously reported in insulin-induced IRE1 phosphorylation by AKT, and subsequent lipogenic upregulation in liver tissue of starved mice^46^. It remains to be elucidated whether *C. elegans* employs a comparable mechanism.

Taken together, by achieving complete rejuvenation in adult organism, we propose that aging-associated decline is not an unidirectional, irreversible process, but a plastic state that can be reset by optimizing the coordination of the proteostasis network.

## Materials and Method

### C. elegans strains

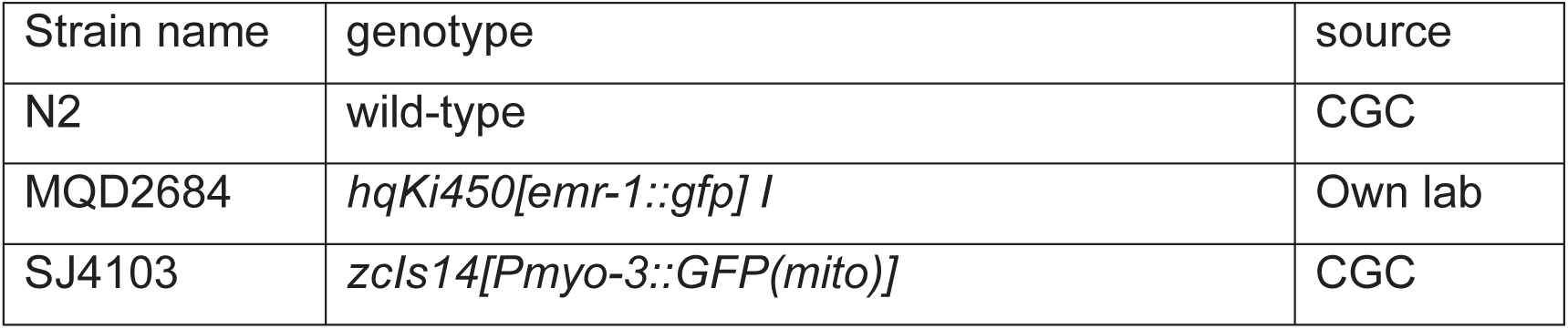
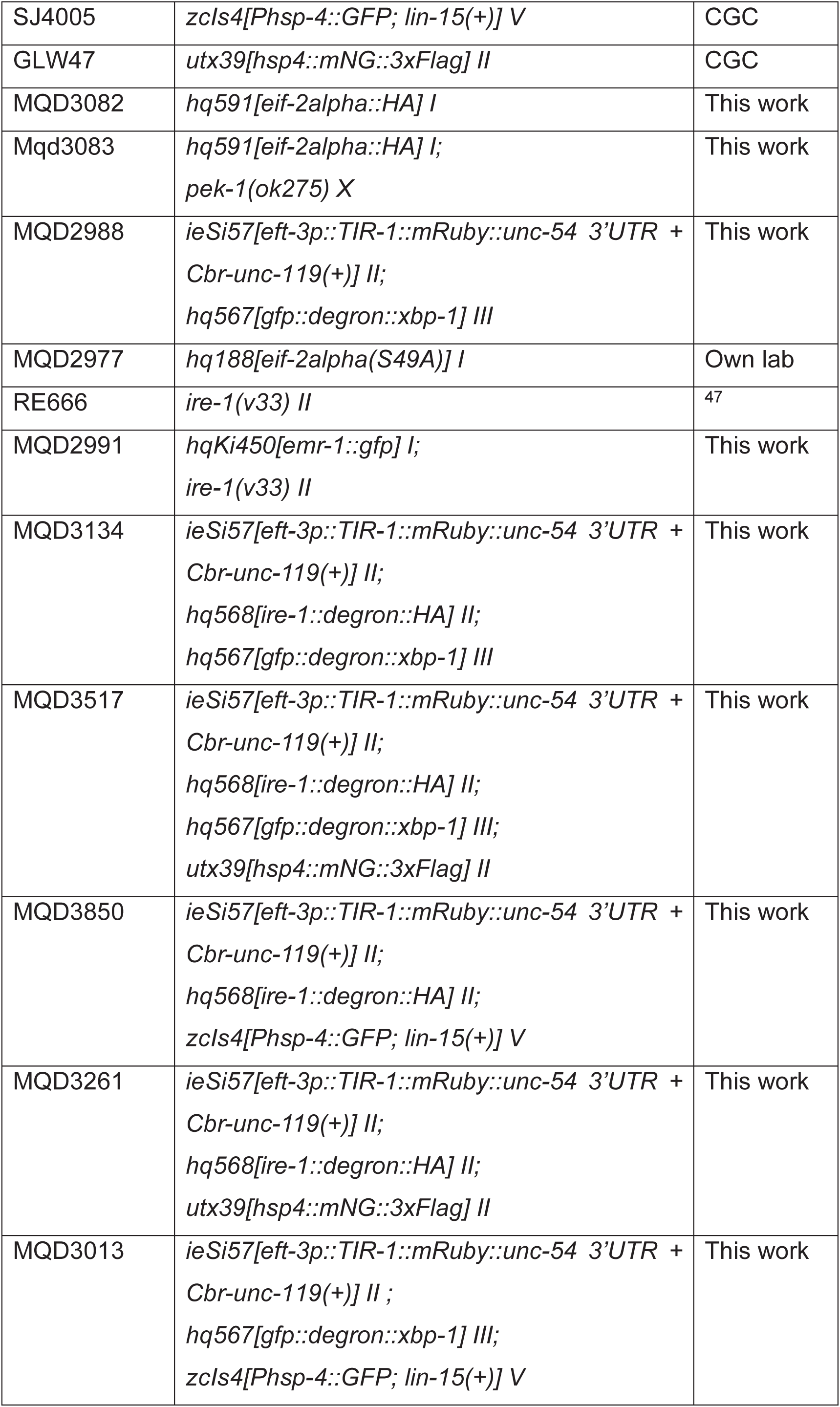
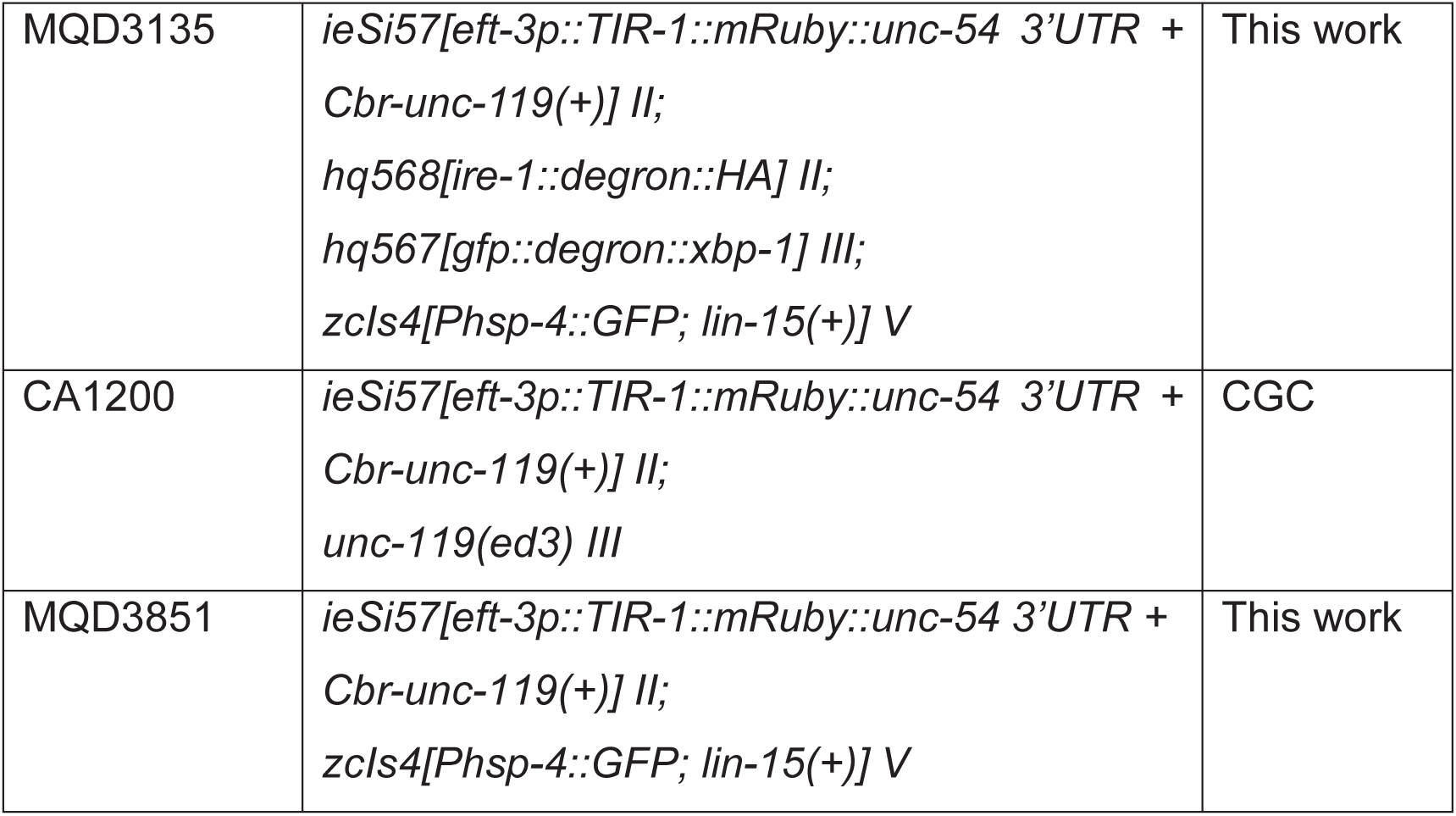

| Strain name | genotype | source |
| --- | --- | --- |
| N2 | wild-type | CGC |
| MQD2684 | *hqKi450[emr-1::gfp]* I | Own lab |
| SJ4103 | *zcls14[Pmyo-3::GFP(mito)]* | CGC |
| SJ4005 | *zcls4[Phsp-4::GFP; lin-15(+)] V* | CGC |
| GLW47 | *utx39[hsp4::mNG::3xFlag] II* | CGC |
| MQD3082 | *hq591[eif-2alpha::HA] I* | This work |
| Mqd3083 | *hq591[eif-2alpha::HA] I;*<br>*pek-1(ok275) X* | This work |
| MQD2988 | *ieSi57[eft-3p::TIR-1::mRuby::unc-54 3'UTR +*<br>*Cbr-unc-119(+)] II;*<br>*hq567[gfp::degron::xbp-1] III* | This work |
| MQD2977 | *hq188[eif-2alpha(S49A)] I* | Own lab |
| RE666 | *ire-1(v33) II* | 47 |
| MQD2991 | *hqKi450[emr-1::gfp] I;*<br>*ire-1(v33) II* | This work |
| MQD3134 | *ieSi57[eft-3p::TIR-1::mRuby::unc-54 3'UTR +*<br>*Cbr-unc-119(+)] II;*<br>*hq568[ire-1::degron::HA] II;*<br>*hq567[gfp::degron::xbp-1] III* | This work |
| MQD3517 | *ieSi57[eft-3p::TIR-1::mRuby::unc-54 3'UTR +*<br>*Cbr-unc-119(+)] II;*<br>*hq568[ire-1::degron::HA] II;*<br>*hq567[gfp::degron::xbp-1] III;*<br>*utx39[hsp4::mNG::3xFlag] II* | This work |
| MQD3850 | *ieSi57[eft-3p::TIR-1::mRuby::unc-54 3'UTR +*<br>*Cbr-unc-119(+)] II;*<br>*hq568[ire-1::degron::HA] II;*<br>*zcls4[Phsp-4::GFP; lin-15(+)] V* | This work |
| MQD3261 | *ieSi57[eft-3p::TIR-1::mRuby::unc-54 3'UTR +*<br>*Cbr-unc-119(+)] II;*<br>*hq568[ire-1::degron::HA] II;*<br>*utx39[hsp4::mNG::3xFlag] II* | This work |
| MQD3013 | *ieSi57[eft-3p::TIR-1::mRuby::unc-54 3'UTR +*<br>*Cbr-unc-119(+)] II ;*<br>*hq567[gfp::degron::xbp-1] III;*<br>*zcls4[Phsp-4::GFP; lin-15(+)] V* | This work |
| MQD3135 | *ieSi57[eft-3p::TIR-1::mRuby::unc-54 3'UTR + Cbr-unc-119(+)] II;*<br>*hq568[ire-1::degron::HA] II;*<br>*hq567[gfp::degron::xbp-1] III;*<br>*zcls4[Phsp-4::GFP; lin-15(+)] V* | This work |
| CA1200 | *ieSi57[eft-3p::TIR-1::mRuby::unc-54 3'UTR + Cbr-unc-119(+)] II;*<br>*unc-119(ed3) III* | CGC |
| MQD3851 | *ieSi57[eft-3p::TIR-1::mRuby::unc-54 3'UTR + Cbr-unc-119(+)] II;*<br>*zcls4[Phsp-4::GFP; lin-15(+)] V* | This work |

### Bacterial strains

*E. coli* OP50 (from CGC)

*E. coli* OP50-expressing GFP (from CGC)

*E. coli* OP50-expressing mCherry (from ^48^)

### Worm maintenance and experimental conditions

Worms were maintained at 20°C, and all experiments were conducted at 25°C. M9 buffer was supplemented with 100 µg/mL ampicillin to reduce the adherence of worms to plastic and prevent bacterial growth.

### Induction of and recovery from adult reproductive diapause

Synchronized worms were allowed to grow until mid-L4 stage on live OP50. The mid L4 stage was determined by vulva development ^49^. For starvation treatment, worms were washed off the plates with M9 buffer. After centrifugation at 3000 rpm for 30s, supernatant was removed, and washing was repeated five times. Then, worms were transferred to petri dish containing M9 buffer for starvation. For refeeding treatment, worms were collected into EP-tubes. After centrifugation at 3000 rpm for 10s, supernatant was removed, and worms were transferred onto NGM plates seeded with *E. coli* OP50.

### Lifespan assay

After 5 or 15 days of starvation, worms were refed on NGM plates seeded with *E. coli* OP50. Post-refeeding lifespans were determined by scoring a population of about 120 worms every other day. Worms that did not respond to a gentle touch were determined as dead. Worms that bagged, showed vulval bursting, or crawled off the plate were censored.

### Imaging and image analysis

Confocal imaging was performed using spinning-disk confocal microscope (UltraVIEW VOX, PerkinElmer, equipped with a 63x, 1.4 numerical aperture oil-immersion objective; or ANDOR Dragonfly 200, equipped with a 60x, 1.4 numerical aperture oil-immersion objective). Worms were placed on 3% agarose pads and anesthetized with 10 mM levamisole solution. The exposure time and laser power were varied to balance the fluorescence intensity among samples. Nuclear morphology in hypodermis and mitochondrial morphology in body wall muscle was manually classified into three classes. Fluorescence intensity was quantified using ImageJ.

### Intestinal barrier function analysis

Worms were transferred into 1.5 mL M9 buffer mixed with 150 µL 11x fluorescein stock solution in a 1.5 mL EP tube. After 3 hours of incubation, worms were washed several times with M9 buffer until no visible color remained in the supernatant. Worms containing fluorescein in intestinal lumen and/or intestine were classified as “non-smurf”, and those containing fluorescein throughout body cavity were classified as “smurf”. Leaky percentage (leaky%) was calculated as “smurf / (smurf + non-smurf) * 100%”.

### Locomotion analysis

To rule out any confounding effects of food preference, locomotion analysis was performed on non-seeded NGM plates ^25^. A population of about 100 worms were transferred onto non-seeded NGM plates and their movement was recorded for 10s. Videos were analyzed by wrMTrck, which is a plugin for ImageJ.

### Western blotting

For SUnSET assay, worms were collected into 900 µL M9 buffer in 1.5 ml tubes. 100 µL 10 mg/mL puromycin stock solution was added to a final concentration of 1 mg/ml. Incubate with rotation for 1 h in the dark. The following antibodies were used: anti-puromycin clone 12D10 (MERCK MABE343, 1:5000 dilution), anti-mouse IRDye 680RD (LICORbio 926-68072, 1:10000 dilution). For eIF2a phosphorylation assay, the following antibodies were used: anti-phospho-eIF2a (Ser51) (Cell Signaling Technology 3398S, 1:5000 dilution), anti-HA (Medical & Biological Laboratories M180-3, 1:10000 dilution), anti-rabbit IRDye 800CW (LICORbio 926-32211, 1:10000 dilution), anti-mouse IRDye 680RD (LICORbio 926-68072, 1:10000 dilution).

### Auxin inducible degradation

20 mM natural auxin indole-3-acetic acid (MedChemExpress, HY-18569) stock solution was prepared freshly (in M9 buffer containing 8% ethanol (v/v)). To prepare IAA plates, IAA stock solution or M9 buffer containing 8% ethanol was added to NGM plates seeded with OP50, to a final concentration of 1 mM IAA or 0.4% ethanol. For IAA pre-treatment, worms were incubated in M9 buffer containing 1 mM IAA or 0.4% ethanol with rotation in dark for 3 hours. Then, worms were refed on IAA plates or ethanol plates in dark.

### Cycloheximide treatment

20 mM cycloheximide (MedChemExpress, HY-12320) stock solution was prepared freshly (in M9 buffer). To prepare CHX plates, CHX stock solution or M9 buffer was added to NGM plates seeded with OP50, to a final concentration of 1 mM CHX. For CHX pre-treatment, worms were incubated in M9 buffer containing 1 mM CHX with rotation in dark for 3 hours. Then, worms were refed on CHX plates in dark.

### Transcriptome analysis

#### Quantification

For RNA-seq, three independent biological replicates were prepared. Raw reads were processed using fastp to obtain clean data. Clean reads were aligned to the WS235 reference genome using HISAT2. Resulting SAM files were converted to BAM format, sorted, and indexed using Samtools. Gene expression levels were quantified with featureCounts using the WS235.97 annotation. Low-abundance genes were filtered by retaining only those with a mean raw counts >10. To enable cross-sample and cross-gene comparisons, raw counts were normalized to Transcripts Per Million (TPM).

#### Characterization of aging-associated gene expression signatures

BiT age analysis was carried out based on source code from ^20^. Counts-per-million normalized RNA-seq reads (CPM) were used as input for BiT age analysis.

To characterize marker genes of aging, we analyzed a set of 327 aging-related signature genes (260 up-regulated and 67 down-regulated during aging) as defined by Fedichev et al ^21^. For each gene, relative expression changes were calculated as fold changes between each time point (Y) and the baseline (X, e.g., pre-refed). Distribution of fold changes was visualized using boxplots, with whiskers representing the 10th to 90th percentiles.

#### Clustering and enrichment analysis

We performed soft clustering analysis using the Mfuzz R package. Genes with significant temporal variation—defined as an absolute log2 fold change >1 at any time point relative to the baseline (refed 0h)—were selected. A total of 11,348 genes met this criterion and were used for subsequent clustering. These genes were grouped into 8 distinct clusters based on the fuzzy c-means algorithm. GO and KEGG enrichment analyses were performed using the clusterProfiler R package.

### Proteome analysis

#### Protein Extraction and Digestion

Worm pellets were cryo-milled (Mixer Mill MM 400, 30 Hz) and lysed in buffer containing 100 mM Tris, 100 mM DTT, and 4% SDS. Protein concentration was determined using the 2-D Quant Kit (GE Healthcare). For quantitative analysis, 200 μg of lysate was mixed 1:1 with 15N-labeled samples. Using the FASP method (30 KD filter), SDS was replaced with 8 M urea, followed by reduction with 50 mM IAA and overnight digestion with trypsin in 50 mM NH4HCO3 at 37°C.

#### LC-MS/MS Analysis

Peptides were analyzed using an EASY-nLC 1000 coupled with an Q Exactive HF mass spectrometer. Separation was performed over a 240-min gradient (0.1% FA in water/acetonitrile) at 600 nL/min using a 50-cm analytical column (150 µm ID, Luna C18 1.8 µm 100 Å resin, Welch Materials). The MS was operated in data-dependent acquisition (DDA) mode (top 15) with resolutions of 60,000 (MS1) and 15,000 (MS2). Fragmentation was achieved via HCD with a normalized collision energy of 27.

#### Quantification

The MS raw data were searched and quantified using pFind 3.1 against the C. elegans proteome database. pQuant was employed to calculate the ratio of light to heavy peptides, with an FDR threshold set at 1% at both the PSM and protein levels.

#### Clustering and enrichment analysis

Proteins with significant temporal variation—defined as an absolute log2 fold change >0.38 at any time point relative to the baseline (refed 0h)—were selected. These genes were grouped into 8 distinct clusters based on the fuzzy c-means algorithm. GO and KEGG enrichment analyses were performed using the clusterProfiler R package.

### Data availability

All sequencing data can be found on the SRA database under accession number PRJNA1420008. Worm strains are available upon request.

---

**Fig S1.**
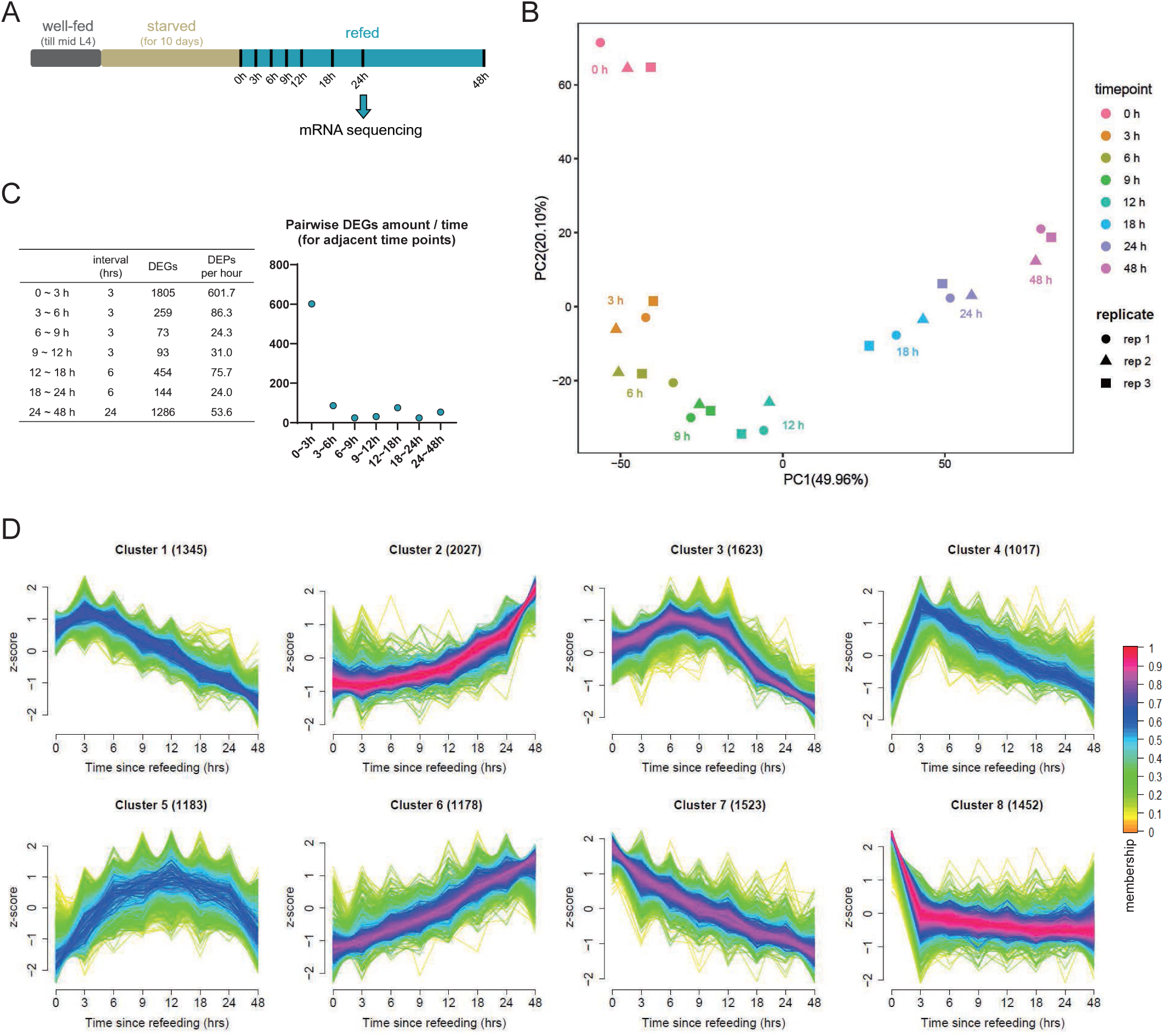
Time-series transcriptomic analysis of refeeding process. (A)Experimental design of time-series transcriptomic analysis. (B)Principal component analysis revealed that refeeding rapidly reshaped transcriptome. (C)Most differentially expressed genes (DEGs) emerged within 3 hours of refeeding. (D)Mfuzz clustering analysis clustered genes by their temporal pattern.

**Fig S2.**
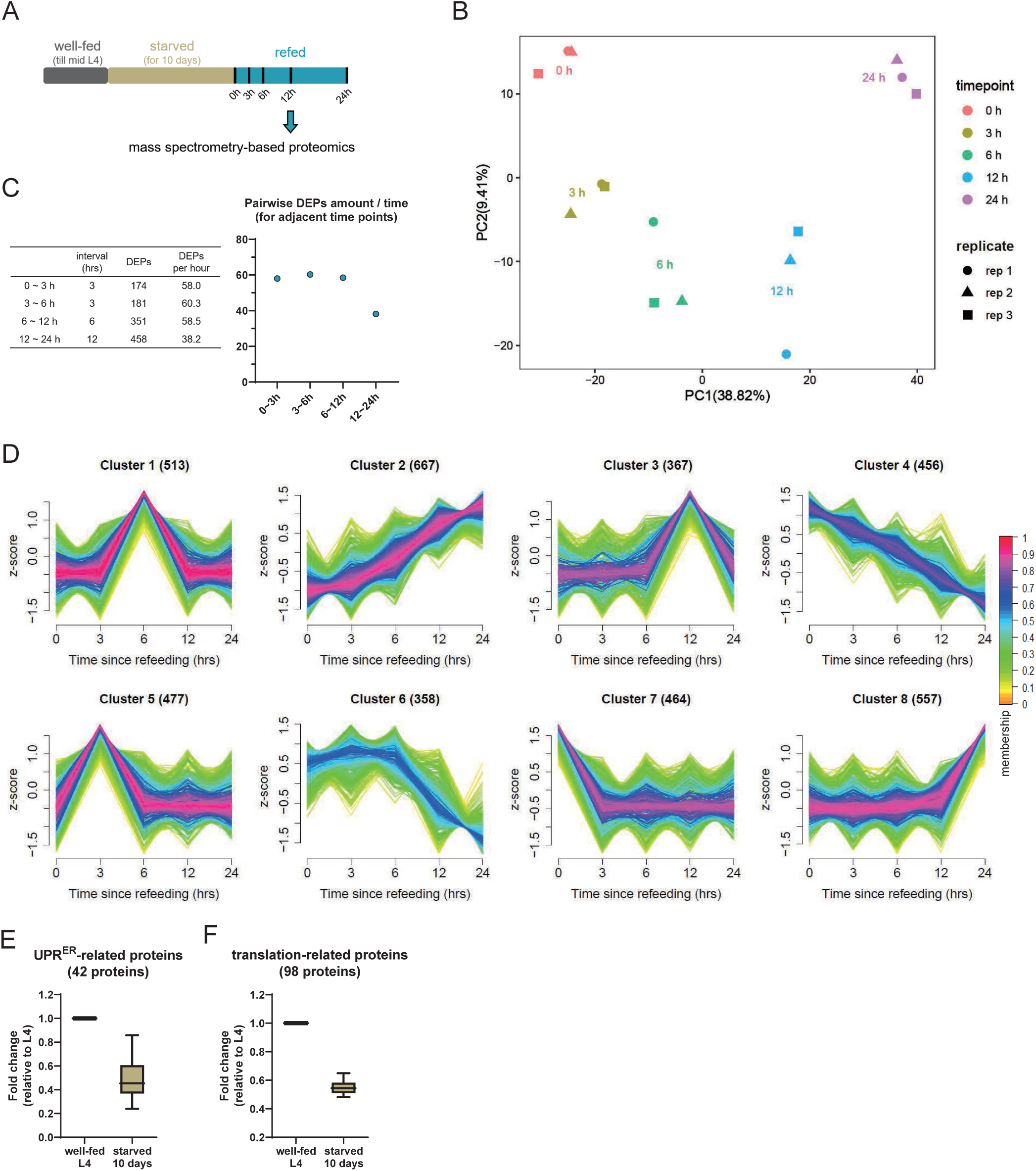
Time-series proteomic analysis of refeeding process. (A)Experimental design of time-series proteomic analysis. (B)Principal component analysis revealed that refeeding gradually reshaped proteome. (C)Differentially expressed proteins (DEPs) gradually emerged after refeeding. (D)Mfuzz clustering analysis clustered proteins by their temporal pattern. (E and F) Abundance of UPR^ER^-related proteins (E) and translation-related proteins (F) decreased during starvation.

**Fig S3.**
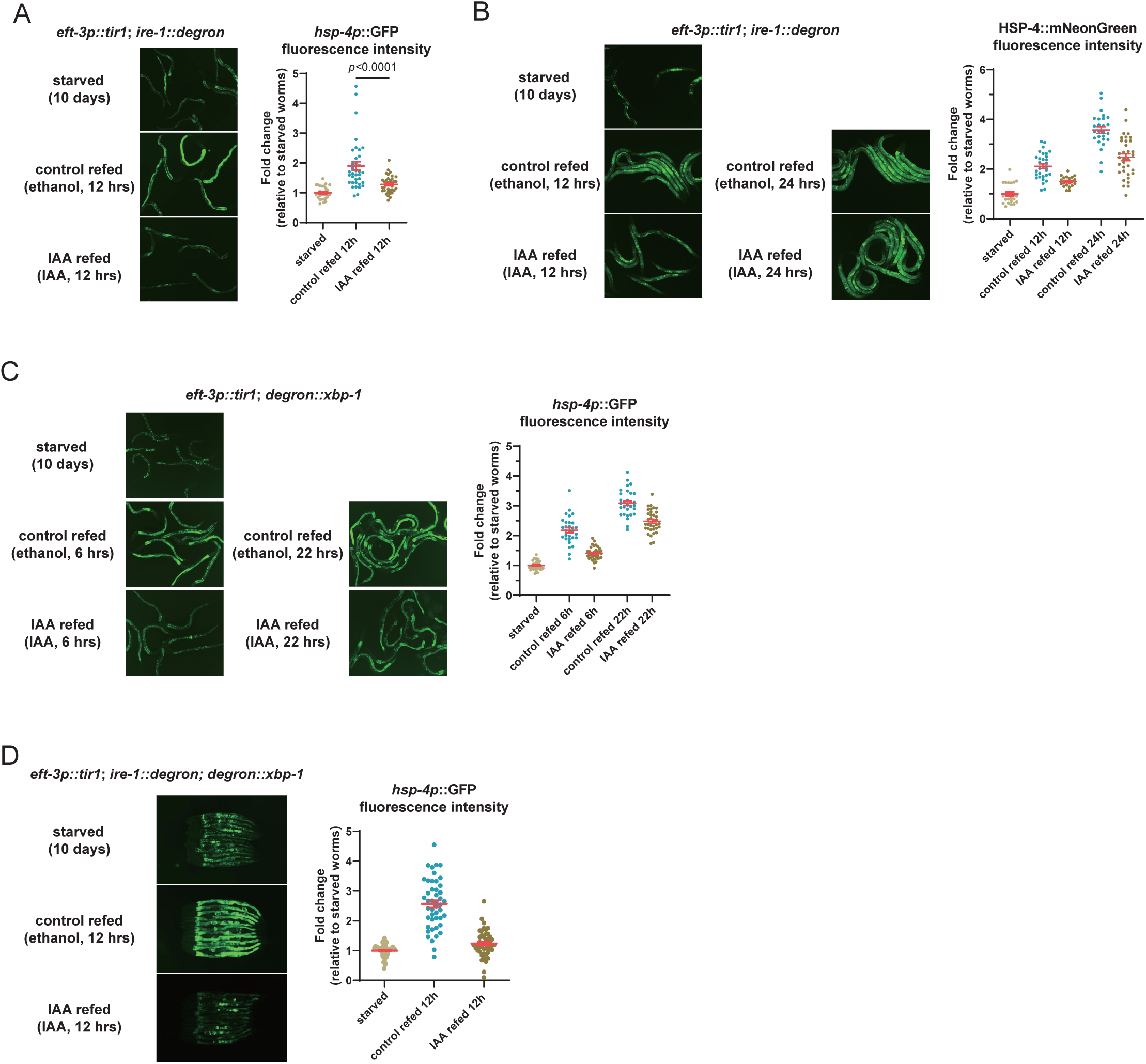
Degradation of IRE-1 or XBP-1 alone partially inhibited UPR^ER^. (A and B) Degradation of IRE-1 largely inhibited transcriptional up-regulation of *hsp-4* (A) but only partially inhibited accumulation of HSP-4 protein. *p*, unpaired Mann-Whitney U test. (c)Degradation of XBP-1 only partially inhibited transcriptional up-regulation of *hsp-4*. (D)Degradation of IRE-1 and XBP-1 largely inhibited transcriptional up-regulation of *hsp-4*. Each dot represents one animal. Error bars are means + SEM.

**Fig S4.**
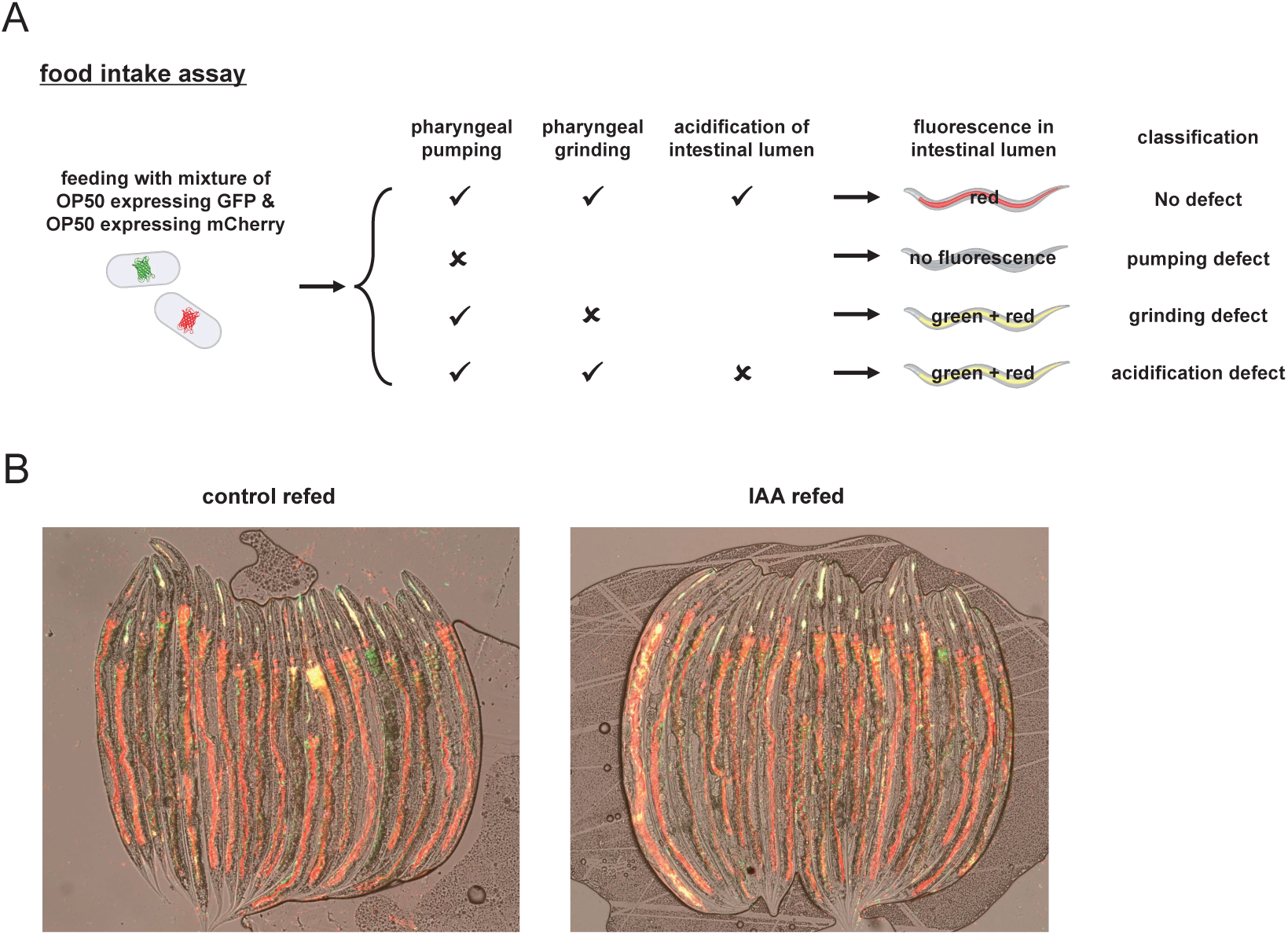
Degradation of IRE-1 and XBP-1 did not lead to defect in food intake. (A)Experimental design of food intake assay. (B)Degradation of IRE-1 and XBP-1 did not lead to defect in food intake.

**Fig S5.**
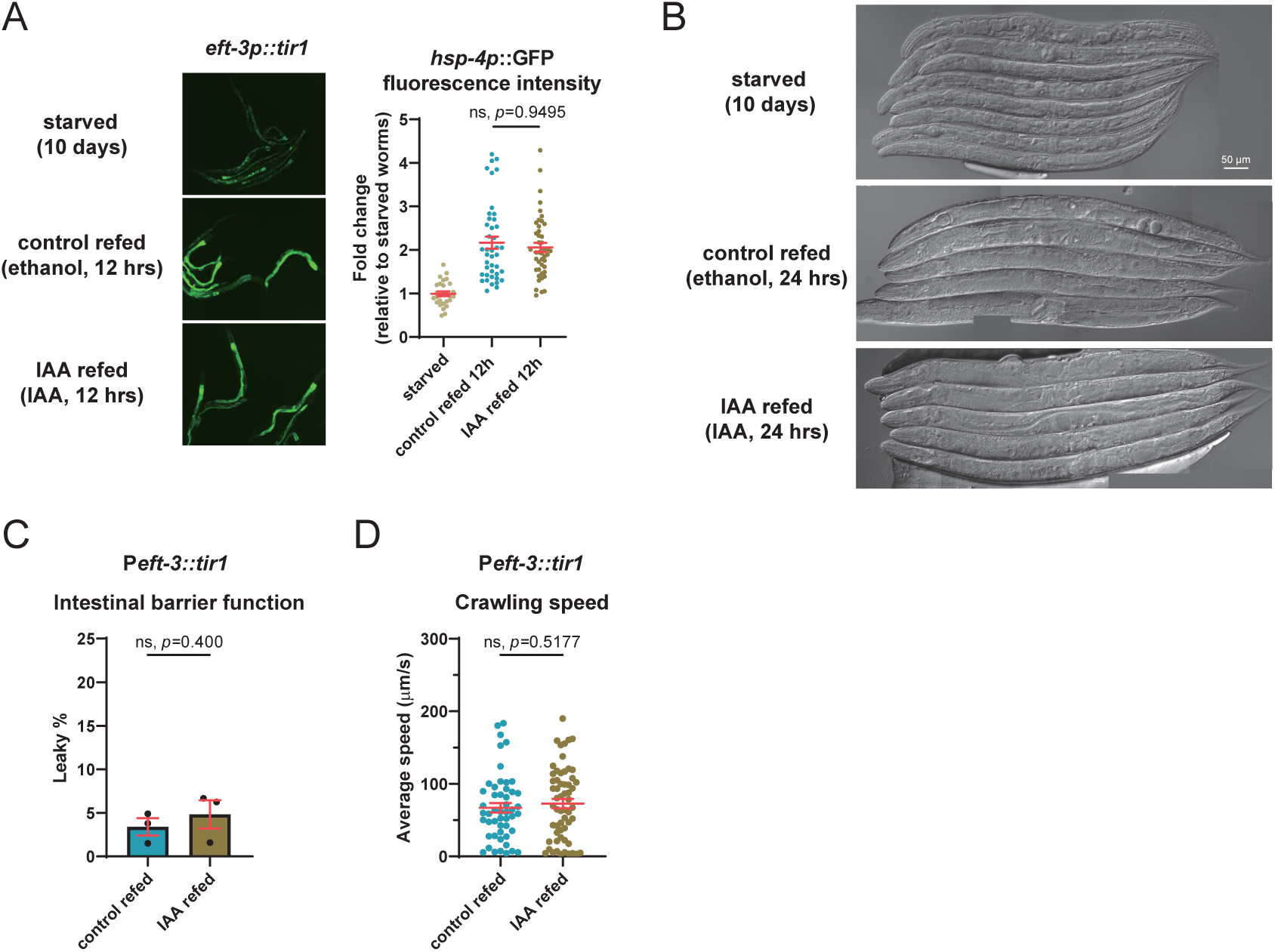
IAA treatment in worms lacking degron tag did not affect refeeding-associated rejuvenation. (A) IAA treatment did not affect transcriptional up-regulation of *hsp-4*. Each dot represents one animal. *p*, unpaired Mann-Whitney U test. (B, C and D) IAA treatment did not affect somatic regrowth (B), reversal of intestinal leaky (C) and restoration of crawling speed (D) in refed worms. In (C), each dot represents one independent biological replicate, *p* represents unpaired t-test. In (D), each dot represents one animal, *p* represents unpaired Mann-Whitney U test. Error bars are means + SEM.

**Fig S6.**
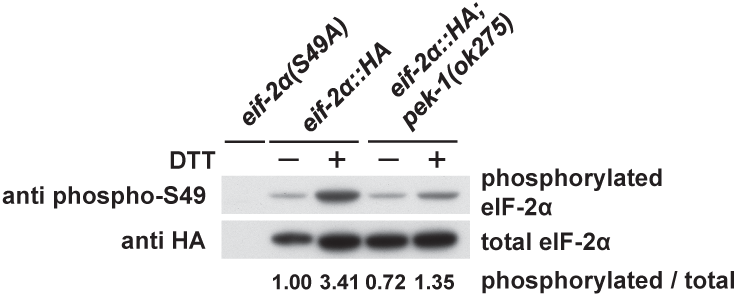
DTT treatment induced phosphorylation of eIF2a. DTT treatment induced phosphorylation of eIF2a. eIF2a(S49A) cannot be phosphorylated at Ser49 site, thus serving as a validation of specificity of the phospho-antibody.

**Fig S7.**
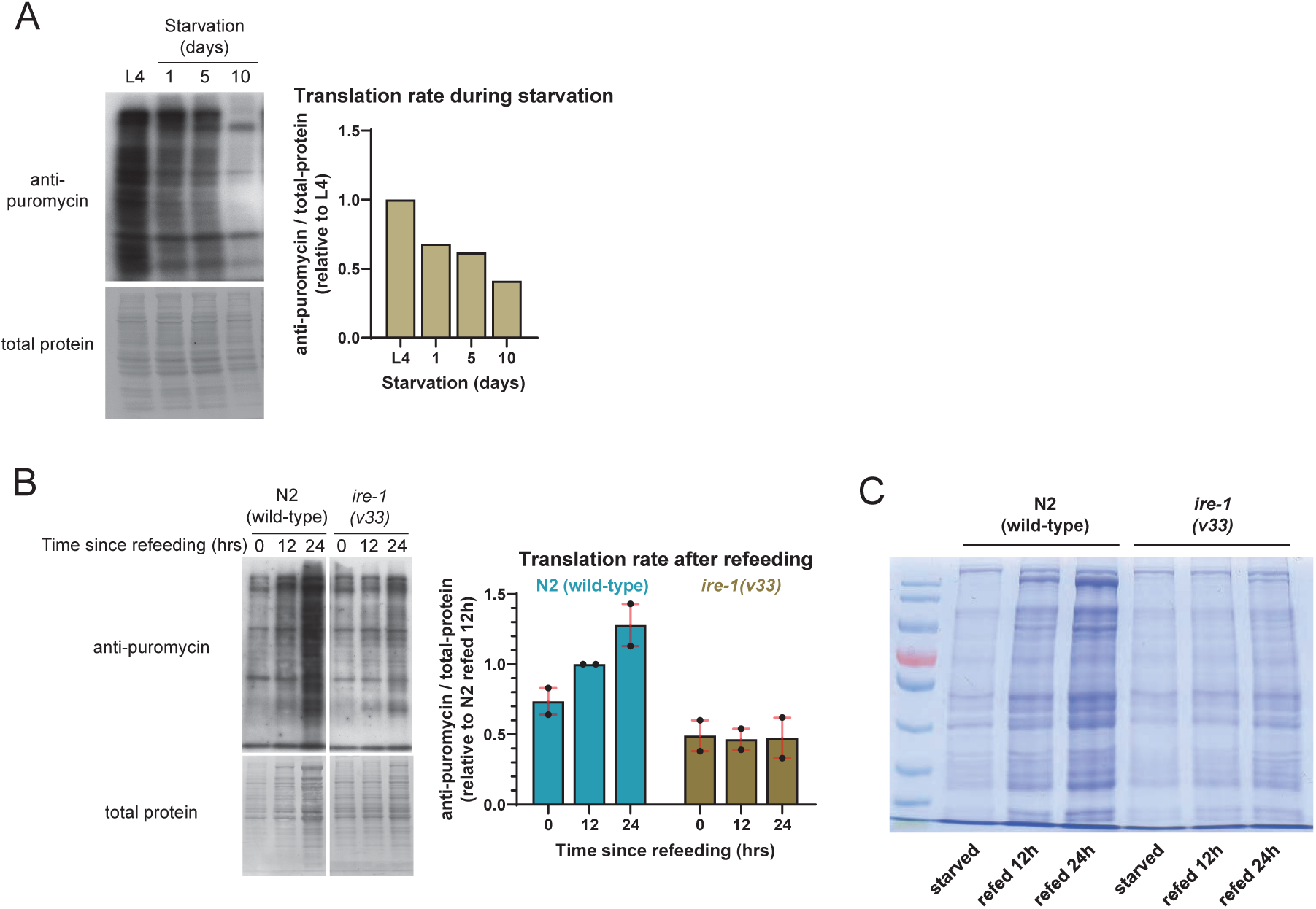
Translation rate decreased during starvation in wild-type worms, and did not increase in refed *ire-1(v33)* mutants. (A)Translation rate (measured by SUnSET assay) decreased during starvation. (B)Translation rate (measured by SUnSET assay) did not increase in refed *ire-1((v33)*. Each dot represents one independent biological replicate. (C)Total protein content did not increase in refed *ire-1(v33)*. Error bars are means + SEM.

**Fig S8.**
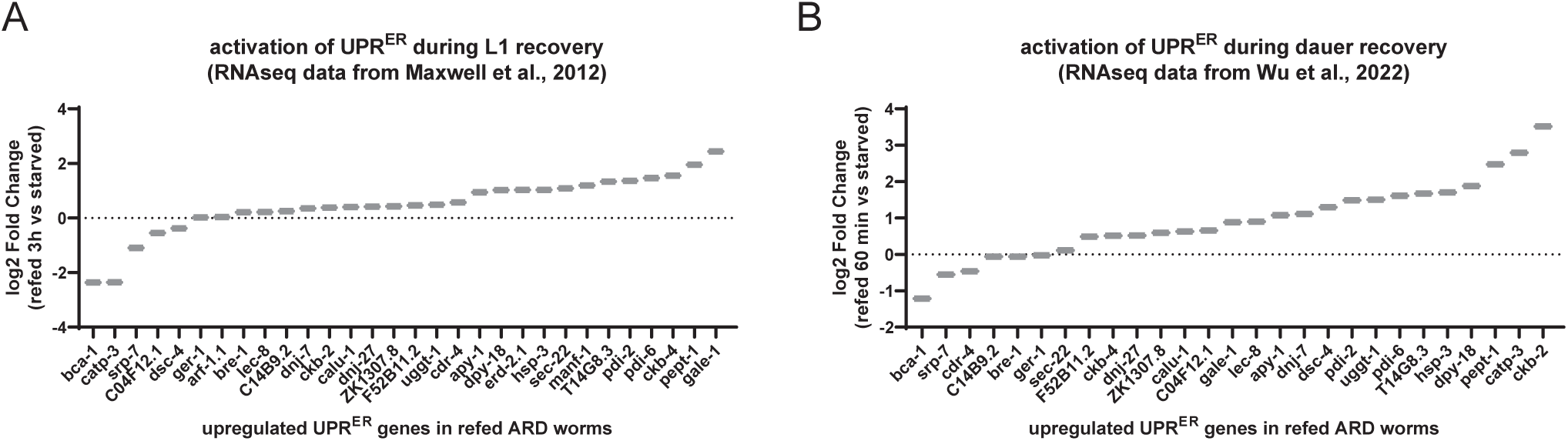
UPR^ER^ activation during L1 recovery and dauer recovery. (A and B) Most UPR^ER^ genes that were activated during ARD recovery were also activated during L1 recovery (A)^37^ and dauer recovery (B)^25^.

**Fig S9.**
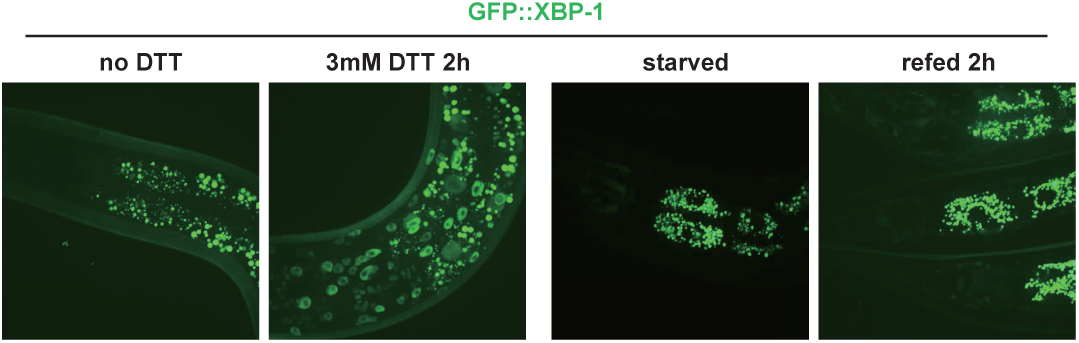
Nuclear translocation of XBP-1 upon DTT treatment, but not in starved or refed worms. Nuclear translocation of GFP::XBP-1 upon DTT treatment but not after refeeding.

**Fig S10.**
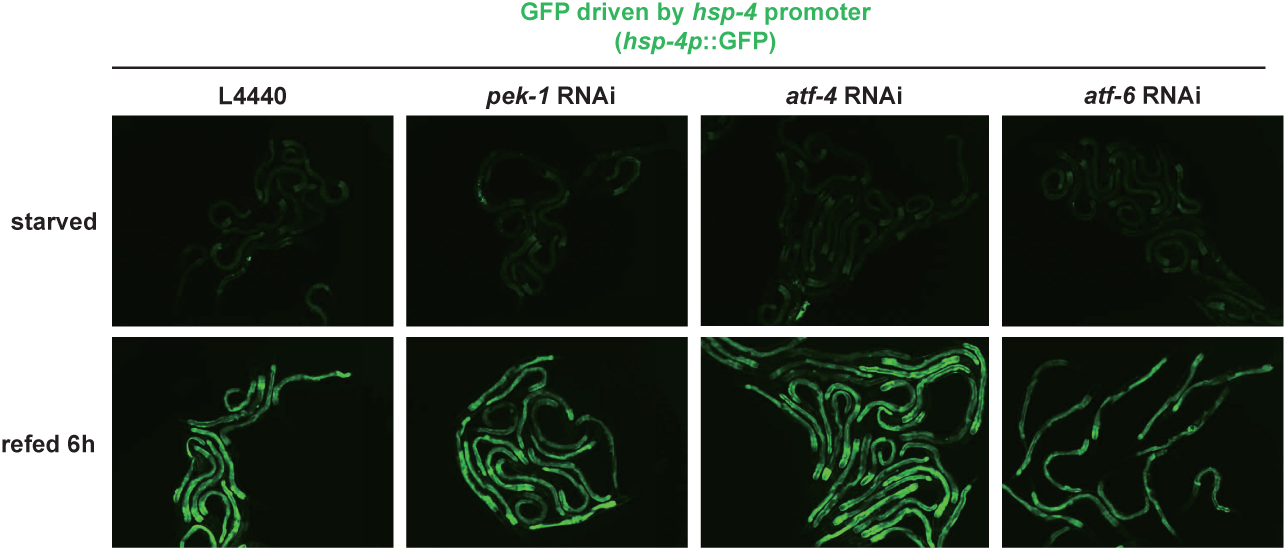
PEK-1 or ATF-6 did not regulate up-regulation of *hsp-4*. Knocking down *pek-1*, *atf-4*, or *atf-6* (since embryo) did not inhibited transcriptional activation of *hsp-4*.

## Notes

### Competing Interest Statement

The authors have declared no competing interest.

